# Environmental Detection of the Amphibian Chytrid Fungus in Water Bodies Predicts Host Infection Along a Deforestation Gradient

**DOI:** 10.1101/2025.08.21.671594

**Authors:** Ananda Brito de Assis, David Rodriguez, Wesley J. Neely, Carla Martins Lopes, Paula Prist, Carlos Arturo Navas, Renato A. Martins, Célio Fernando Baptista Haddad, C. Guilherme Becker

## Abstract

Understanding pathogen dynamics during environmental life stages is vital for comprehending wildlife diseases, especially for those with a free-living phase. In amphibians affected by the chytrid fungus *Batrachochytrium dendrobatidis* (*Bd*), most studies have focused on host-pathogen interactions, with less emphasis on *Bd*’s environmental stage. We tested whether the distribution of *Bd* in natural aquatic environments can predict host infection patterns. We sampled four tropical amphibian species across eight rainforest landscapes with varying habitat loss, testing whether environmental and host *Bd* detection varied along gradients of habitat change. Using a high-capacity water filtration method coupled with digital and real-time PCR detection assays, we quantified *Bd* in water and amphibian samples. Our results revealed a strong positive correlation between *Bd* DNA concentrations in water samples and infection loads on amphibian skin samples. Forest cover and habitat split were the primary predictors of *Bd* distribution in both free-living and host-associated forms. We identified *Bd*-GPL and *Bd*-Asia-2/Brazil lineages across our study landscapes. Our study introduces and validates a robust protocol for detecting *Bd* in environmental samples, with the potential to enhance monitoring and inform management strategies. Moreover, our work contributes novel, well-replicated spatial data on *Bd* associations between hosts and their environments.

## 1. Introduction

The ecology of infectious diseases caused by pathogens persisting in the environment can be particularly complex, primarily because the heterogenous environment itself serves as a reservoir (Rees et al., 2021). Therefore, understanding the biology of pathogens during their environmental or saprophytic stages is a critical aspect in tracking and managing wildlife disease dynamics. Environmental factors can independently or synergistically influence disease dynamics by altering pathogen behavior and survival during their free-living stage and consequently shape host-pathogen interactions (Hoyt et al., 2023; Suh et al., 2024). Chytridiomycosis in amphibians provides a suitable system for studying the dynamics of an infectious disease at the landscape and ecosystem levels. This disease is caused by the amphibian chytrid fungus *Batrachochytrium dendrobatidis* (hereafter *Bd*), a particularly important wildlife pathogen that impacts hundreds of amphibian species globally (Scheele et al., 2019). This fungus is characterized by both a motile, waterborne zoospore stage and a sometimes host-associated sessile zoosporangia stage (Longcore et al., 1999). While substantial research has investigated various aspects of *Bd* during the host-associated stage, relatively little is known about persistence and transmission of this pathogen through environmental reservoirs.

Evidence from laboratory (Gass et al., 2024) and field (Alvarado-Rybak et al., 2021; Rohr and Raffel, 2010) research indicates that *Bd* thrives in microclimates with mild temperatures and high humidity (Turner et al., 2021), like those found in natural, close-canopy montane tropical forests (Becker and Zamudio, 2011). However, the difficulty in detecting and quantifying *Bd* zoospores in natural environments, using environmental DNA methods (*i.e*., DNA extracted directly from environmental samples without prior isolation of any target organism; Taberlet et al., 2012) has hindered researchers from fully comprehending crucial aspects of *Bd* ecology, including its distribution and persistence across diverse habitats (Brannelly et al., 2020; Mosher et al., 2018). The non-detection of *Bd* in environmental samples could stem from several factors, including: (1) limitations in polymerase chain reaction (PCR) sensitivity, where the assay fails to detect low *Bd* concentrations, (2) challenges with filtration methods, as water samples from natural environments can quickly clog filters, limiting filtration to very small volumes, (3) rapid DNA degradation in the environment, especially in warm tropical regions, and (4) interactions among these factors. Environmental DNA (eDNA) in aquatic environments tends to degrade faster at higher temperatures and UV radiation levels (Pilliod et al., 2014; Strickler et al., 2015), which are typical conditions of open-canopy environments. Thus, sampling and detecting environmental *Bd* across a range of ecological conditions is important to elucidate the mechanisms underlying disease transmission and the complex interactions between *Bd* and its habitat.

Anthropogenic habitat modifications can significantly alter the microclimatic profile of remnants of natural habitats (*e.g*., forest fragments) and influence other abiotic and biotic factors such as temperature, humidity, UV radiation, and wind (Fischer et al., 2007). Previous studies have linked habitat degradation to the emergence and outbreaks of a wide variety of infectious diseases (Brearley et al., 2013). However, the complex interactions between environmental disturbances, *Bd* dynamics in hosts, and environmental reservoirs that drive the decline of amphibian populations are still not fully understood. Degraded, open-canopy environments can lead to reduced *Bd* prevalence in amphibian populations due to temperature and humidity constraints (Becker and Zamudio, 2011), although the risk of *Bd* infection appears to be higher in amphibian populations from small, isolated forest fragments compared to continuous, natural forests (Belasen et al., 2022). This suggests that forest fragmentation could be exacerbating disease-induced population declines in amphibians. Furthermore, the role of landscape-scale habitat loss and local-scale habitat fragmentation and split (*i.e*., artificial discontinuity of the multiple habitats types needed by different amphibian life history stages) should impact both host-*Bd* interactions (Becker et al., 2023), as well as the viability of *Bd* in environmental reservoirs. Thus, understanding the drivers of landscape-level environmental, *Bd* persistence would advance both theoretical and applied aspects of amphibian disease ecology, as this could inform *Bd* surveillance from environmental samples when proper host screening is challenging.

In this study, we hypothesized that *Bd* occurrence and concentrations in environmental samples would mirror occurrence and infection intensity patterns (*i.e*., loads) observed in amphibian hosts across different habitats. By examining these factors across four tropical amphibian species sampled along eight landscapes with differing levels of habitat loss and fragmentation, we aimed to environmentally detect *Bd* and then estimate the direct impact of habitat change on the interactions between environmental *Bd* concentration and host-associated *Bd* infection loads. We also leveraged our dataset to compare the sensitivity of digital (dPCR) and quantitative polymerase chain reaction (qPCR) techniques for detecting and quantifying *Bd* DNA targets. Our study provides valuable insights into the applications of these methods for monitoring multiple lineages of *Bd* in environmental samples. More broadly, our findings clarify the role of habitat alteration in driving environmental *Bd* presence which will inform future surveillance efforts of invasive facultative fungal pathogens.

## 2. Materials and Methods

### 2.1. Sampling design

We conducted a survey across eight circular landscapes (10-km radius) within the Southeastern Atlantic Forest of Brazil, encompassing a gradient of deforestation ranging from mostly continuous forest (N = 4 landscapes) to heavily fragmented forests due to agriculture (N = 4 landscapes). Within each focal landscape, we sampled five natural (*i.e*., not silviculture) forest fragments or five continuous forest sites, resulting in a total of 40 sampling sites. Sites were all located within a 10 km radius within each landscape (Figure 1). Sampling for *Bd* was conducted at two levels: environmental (stream water filtering) and host (amphibian skin swab) pathogen reservoirs (Table SI 1).

**Figure 1.**
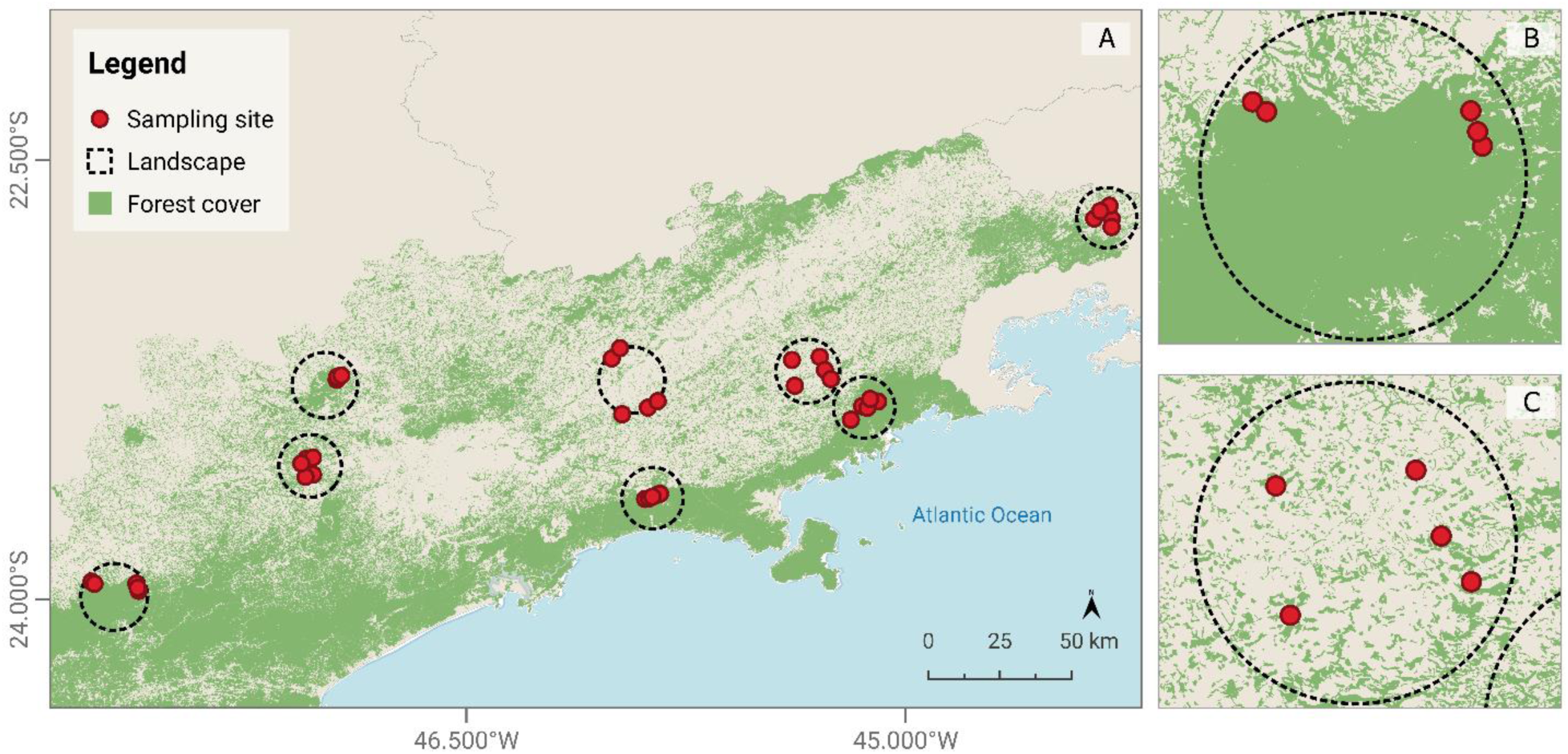
Sampling Area. (A) The eight focal landscapes within the Southeastern Atlantic Forest of Brazil, with forest cover shown in green. Each dashed black circle represents a 10-km radius landscape, with five red dots indicating sampling sites. (B) A magnified section from (A) illustrating an example of a continuous forest landscape. (C) A magnified section from (A) illustrating an example of a fragmented forest landscape.

In each landscape, we sampled five local streams using water filtering techniques and collected a total of 40 water samples. We sampled amphibians alongside the streams and terrestrial environments at each site focusing on four abundant species at the sampling sites: *Ischnocnema henselii* (Brachycephalidae), *Rhinella ornata* (Bufonidae), *Boana faber* (Hylidae), and *Haddadus binotatus* (Craugastoridae). The number of amphibians sampled per landscape depended on availability at the time of collection but typically exceeded 10 individuals (Table SI 1). All sampled individuals were immediately released at the capture location after processing through non-invasive skin swabbing. Host and environmental sampling occurred during the latter portion of the amphibian breeding season (Bertoluci, 1998), from January 20 to March 28, 2022.

### 2.2. Landcover metrics

Spatial data at the landscape and site levels, including hydrological features and land use/land cover, were obtained from the repository of the Brazilian Foundation for Sustainable Development (Fundação Brasileira Para o Desenvolvimento Sustentável). Land cover layers generated using RapidEye satellite imagery at a 1:10,000 scale, had a resolution of 5 x 5 meters and corresponded to the year 2017 (Fundação Brasileira Para o Desenvolvimento Sustentável). These were the available data at the time of sampling. These data encompassed six land cover classifications including both natural habitats as well as anthropogenic land cover such as agriculture, silviculture, and urbanization, along with drainage networks comprising streams and ponds (small permanent water bodies were mapped by visual classification in Google Earth and in ArcGIS and integrated into this dataset). From this dataset, we derived 18 environmental metrics, measured at the landscape (10 km radius) and local scales (250 m radius), considered both land cover and landscape configuration: forest cover, drainage percentage, edge density, habitat split (*i.e.*, spatial discontinuity between natural forest vegetation and aquatic breeding sites such as streams and ponds) and connectivity, see Supplementary Material and Table SI 2 for more detail. By examining variation in habitat split among forest fragments with comparable sizes and edge densities, we attempted to disentangle the effects of forest fragmentation from those of habitat split.

### 2.3. Water filtering

We conducted water sampling by directly filtering from focal streams using high-capacity filtration capsules (Envirocheck HV^®^) with 1 μm pore-size. This pore diameter is suitable for collecting our target *Bd* zoospores that range between 3-5 μm in diameter (Longcore et al., 1999). We conducted water pumping using a peristaltic pump (model 140, Solinst Canada Ltd., Georgetown, Ontario, Canada), applying a flow rate of 1.5 L/min. We used new piping to connect each capsule to streams, and wore disposable gloves to prevent cross-contamination.

After filtering, we filled capsules with 150 ml of a DNA stabilizing reagent (Tris-HCl 0.1 M, EDTA 0.1 M, NaCl 0.01 M, and N-lauroyl sarcosine 1%, pH 7.5–8) to prevent DNA degradation (Lopes et al., 2017). We stored samples at room temperature for a maximum of three days and then transferred them to a freezer at −20°C until DNA extraction. At each sampling stream, we filtered the maximum volume of capsule’s capacity, resulting in a mean of 94.6 L (+/− 57.8 L) of water per water body. We took a negative control sample by filtering 5 liters of distilled water to monitor for potential contamination of equipment. This negative control underwent the same storage, transport, extraction, and PCR procedures as the environmental samples, ensuring that we monitored potential contamination throughout all procedural steps.

### 2.4. *Bd* screening in amphibian hosts

We screened amphibians for *Bd* immediately after stream water sampling. Animals were captured through active searches at night using fresh, disposable nitrile gloves for each individual. In total, we sampled 271 individuals assigned to four species: *Boana faber* (N = 104), *I. henselii* (N =69), *H. binotatus* (N = 66), and *R. ornata* (N = 32). We collected skin-swab samples from each frog by gently rubbing the swab against dorsal and ventral surfaces along the rostro-cloacal length five times, and five times on forelimbs and hindlimbs. Swabs were then preserved frozen in dry tubes until DNA extractions and animals were immediately released at the capture location.

### 2.5. DNA extraction

We used the commercial column-based kit NucleoSpin® Soil (Macherey-Nagel, Düren, Germany) to extract total DNA from the stream water samples and amphibian skin swabs. For water samples, we performed additional steps before following the kit manufacturer’s protocol. We adapted the procedure described by Pont et al. (2018), where each filtration capsule filled with 150 ml of buffer solution was agitated for 30 minutes at 250 rpm, then split into 50 ml centrifuge tubes and centrifuged for 20 minutes at 15,000 x g at 6°C. Most of the supernatant was removed and 33 ml of absolute ethanol and 1.5 ml of 3 M sodium acetate were added to each tube containing the pellet. The tubes were incubated overnight at −20°C. After this period, tubes were centrifuged again and the supernatant was discarded. We then added 720 µL of ATL buffer from the kit DNeasy Blood & Tissue Extraction Kit (Qiagen) and, after vortexing, we added 20 μL of Proteinase K. All tubes were then incubated for 2 hours at 56°C. After this step, we proceeded with the NucleoSpin® Soil kit protocol, following the manufacturer’s recommendations starting from step 6 of the manual. We incorporated negative extraction controls, for each 36 samples, comprising solely of extraction reagents, without any sample addition, to monitor potential cross-contamination during the extraction process.

### 2.6. dPCR amplification in water samples

We prepared the dPCR reaction mix with 2.0 µL of Absolute Q™ DNA Digital PCR master mix (2X, Applied Biosystems), 0.5 µL of the SNP assay mix (20X), 4.5 µL of nuclease-free water, and 3 µL of the DNA template in a total volume of 9 µL. The 20X assay comprised of primers ITS1-3 Chytr (18 μM) and 5.8S Chytr (18 μM), along with the Chytr MGB2 probe (8 µM) (Applied Biosystems) (Boyle et al., 2004). We loaded the dPCR reaction mix onto QuantStudio™ MAP16 plates (ThermoFisher Scientific), then added 15 μL of isolation buffer (ThermoFisher Scientific) to the wells over the dPCR mix. We performed acquisition and analysis using Applied Biosystems QuantStudio Absolute Q Digital PCR system (Applied Biosystems). The thermocycling conditions were a preheat at 95°C for 10 min, 40 cycles of 96°C for 5 s followed by 60°C for 15 s. We ran one no-template control along with our samples to monitor for contamination during preparation.

### 2.7. qPCR amplification in host and water samples

We performed real-time quantitative polymerase chain reaction (qPCR) for the quantitative detection of *Bd*. We used real-time TaqMan PCR assay to detect and quantify *Bd* in both water and amphibian samples (Boyle et al., 2004). The assays were performed using the QuantStudio™ 3 Real-Time PCR system (Applied Biosystems). Each sample was analyzed in duplicates using an internal positive control (IPC, Garland et al., 2010). We included two PCR negative controls in each 96-well assay plate using molecular grade water. For the quantification of *Bd* in amphibian samples, we utilized gBlocks Gene Fragments synthetic standards (IDT Technologies) with 10x serial dilutions ranging from 1.95 × 10^2^ to 10^6^ gene copies per microliter (gc/μL). Water assays employed *BdBsal* standards (Pisces Molecular, Boulder, CO) with concentrations ranging from 1.3 × 10^2^ to 1.3 × 10^6^ gc/μL. A third run was conducted when results did not match between duplicates.

The data obtained by using the dPCR and qPCR methods were analyzed using the following *Bd* metrics: (1) *Bd* occurrence (presence/absence) in stream water samples, (2) *Bd* occurrence in amphibian skin samples, (3) *Bd* concentration in water samples (*Bd* DNA copies/L), (4) *Bd* infection loads in host samples (*Bd* DNA copies per whole skin swab).

### 2.8. *Bd* genotyping

For *Bd* genotyping, we employed mitochondrial and nuclear SNP-based assays to differentiate between the global panzootic lineage (*Bd*-GPL), the Brazilian lineage (*Bd*-Asia2/Brazil), and a hybrid lineage (Carvalho et al., 2023; Jenkinson et al., 2018). We only conducted genotyping for *Bd*-positive amphibian skin swabs as these samples contained higher concentrations of *Bd* and thus had a higher likelihood of genotyping success (Neely et al., 2025). We used the following recipe per sample: 5.0 µL TaqMan Fast Advanced Master Mix (ThermoFisher), 0.5 µL 20X SNP Assay (either Bdmt_26360 or BdSC9_621917; reporters 1 and 2 at 4 µM and forward and reverse primers at 18 µM), 1.0 µL nuclease-free water, and 2.5 µL of diluted template DNA. We ran genotyping qPCRs on a QuantStudio™ 5 Real-Time PCR system (Applied Biosystems) using the following cycling protocol: 60 °C for 30 s, then 95 °C for 20 s, then 50 cycles of 95 °C for 1 s and 60 °C for 20 s, then 60 °C for 30 s. We included the following positive controls: Bd-GPL (NAF-01), a hybrid strain (CLFT-024-2), and Bd-Brazil (BAF-038). Molecular grade water was used as the negative control. Genotype calling followed procedures described by Carvalho et al. (2023).

### 2.9. Statistical analysis

#### 2.9.1. Comparing dPCR and qPCR methods

We compared both dPCR and qPCR methods for their effectiveness in detecting and quantifying *Bd* from stream water samples. To examine the level of agreement between the two PCR techniques in detecting *Bd* DNA in water samples, we employed Cohen’s kappa statistics. We also used Spearman’s rank correlation to compare *Bd* concentration between the two methods.

#### 2.9.2. Correlations

We performed Spearman’s rank correlation to test for an association between *Bd* concentration in environmental samples and *Bd* infection load in amphibian skin throughout our sampling sites. For this analysis, we used the log-transformed *Bd* concentration (*Bd* DNA copies/L) from each of the 40 streams across all sampling sites and calculated the average for each landscape (N = 8), and log-transformed *Bd* infection load (total *Bd* DNA copies) per individual frogs, across all species, by sampling sites, and calculated the average for each landscape (N = 8).

#### 2.9.3. Comparing Bd parameters among habitats

We first analyzed the effect of habitat structure on the distribution of *Bd* by comparing two categories of landscape integrity: predominantly continuous (> 50% of natural forest cover) and heavily fragmented forests (<50% of natural forest cover). To conduct this analysis, we employed a combination of Chi-squared tests and Wilcoxon rank-sum tests. Data from both dPCR (water samples) and qPCR (amphibian skin swab samples) techniques were included in the analysis. The Chi-squared test tested whether there was a significant difference in the occurrence (presence/absence) of *Bd* in continuous versus fragmented forests. The non-parametric Wilcoxon rank-sum test was used to compare *Bd* concentration (water samples) or infection loads (host samples) between these forest types, as residuals from Gaussian models did not meet the normality assumptions required for parametric tests. We conducted all analyses related to the selection of land cover metrics using R version 4.4.0 (R Core Team, 2024).

#### 2.9.4. Choice of land cover metrics

We selected land cover metrics to be included as explanatory variables in our analyses using a systematic screening method. The first step involved a visual inspection of a Principal Component Analysis (PCA) biplot using all 18 metrics (see Table SI 2 for the full set) to identify variables with strong contributions to the principal components and to exclude those exhibiting high cross-correlations. The following variables were selected: drainage percentage, forest edge density (forest fragments and riparian protected areas), and percentage of natural forest cover, measured at local (250 m radius) and landscape (10,000 m radius) scales. Habitat split and forest patch connectivity were measured at local and landscape scales, respectively. These variables were included in an AICc model selection pipeline (see below for details), and additional variables were pruned after we detected significant collinearity by calculating Variance Inflation Factor (VIF) for variables in resulting models. Variables with VIF values > 4 were systematically evaluated (for their importance as determinants of *Bd* occurrence, concentration, or load) and excluded from models to reduce multicollinearity. Following these steps, our final list of local scale explanatory variables included: percentage of drainage, forest edge density, natural forest cover, and habitat split. At the landscape scale, the following variables were included in downstream model selection: percentage of drainage in riparian protected area, forest edge density, natural forest cover, and forest patch isolation. Amphibian species and landscape were also included in the model as a fixed effect.

#### 2.9.5. Multivariate Models Linking Habitat Structure and Bd from host and water samples

We built Generalized Linear Mixed Models (GLMMs) to examine the relationships between habitat structure metrics and *Bd* concentration in the water samples, as well as *Bd* infection load in amphibian skin samples. Given that our response variables were count-based (*Bd* DNA copy numbers) and exhibited overdispersion along with an excess of zero values (absence of *Bd* detection in many samples), we modeled the data using a zero-inflated negative binomial distribution (Brooks et al., 2022). Additionally, because samples were nested within landscapes, and thus potentially non-independent, we included landscape as a random effect to account for this hierarchical structure. For modeling *Bd* occurrence (presence/absence), we employed a binomial GLMM. All models were built using the glmmTMB package and ‘logit’ link function (Brooks et al., 2022), which accommodates complex data distributions and random effects while allowing efficient handling of zero-inflated outcomes. We analyzed data at the local level and the landscape level separately, then we built models that integrated the most significant explanatory variables from both scales. We used Moran’s I test to confirm spatial independence of residuals and included landscape as a random effect in all models. We ran model diagnostics through residual analysis, inspecting the dispersion, distribution of residuals, and variance, using the DHARMa package in R (Hartig and Hartig, 2017). We applied Akaike Information Criterion (AICc) model selection to rank all possible models, considering those models with low AIC values, and ΔAIC ≤ 2, with substantial support (Burnham and Anderson, 2002). We used R version 4.4.0 for all analyses (R Core Team, 2024).

## 3. Results

We detected the presence of *Bd* in 23 out of 40 (57.5%) stream water samples and 59 out of 271 sampled amphibian individuals. Specifically, *Bd* was found in 41 of 69 *I. henselii*, 6 of 32 *R. ornata*, 12 of 104 *B. faber*, and 0 of 66 *H. binotatus*. Negative controls did not exhibit amplification throughout the analysis. Furthermore, a zero-inflated negative binomial model revealed no significant effect of water volume on the estimated *Bd* DNA concentration, as determined by both dPCR (Estimate = - 0.001, *p* = 0.917) and qPCR (Estimate = - 0.001, *p* = 0.793).

### 3.1. Comparing techniques for detecting and quantifying *Bd*

We found a strong positive correlation between the *Bd* DNA concentration in water samples when comparing dPCR and qPCR techniques (Spearman, ρ = 0.813; *p* < 0.001). Using dPCR, we detected *Bd* DNA in 47.5% (19/40) of water samples, while with qPCR we detected *Bd* DNA in 57.5% (23/40). Despite this difference in *Bd* DNA detection rates, there was a strong agreement between the two techniques (Cohen’s kappa, k = 0.630, z = 3.89, *p* < 0.001), with a total of 80% (32/40) of samples showing concordant results. Therefore, for subsequent analyses of *Bd* in water samples, we focus on data on *Bd* concentration and occurrence obtained by the dPCR technique, which provides absolute quantification of *Bd* concentration (*Bd* DNA copies/L) per sample.

### 3.2. Environmental *Bd* as a predictor of infection patterns in amphibian hosts

We found a positive correlation between environmental *Bd* DNA concentration in stream water samples and *Bd* infection load in our focal amphibian species (Spearman’s ρ = 0.496, N = 8, *p* < 0.004, Figure 2) for all sampled landscapes.

**Figure 2.**
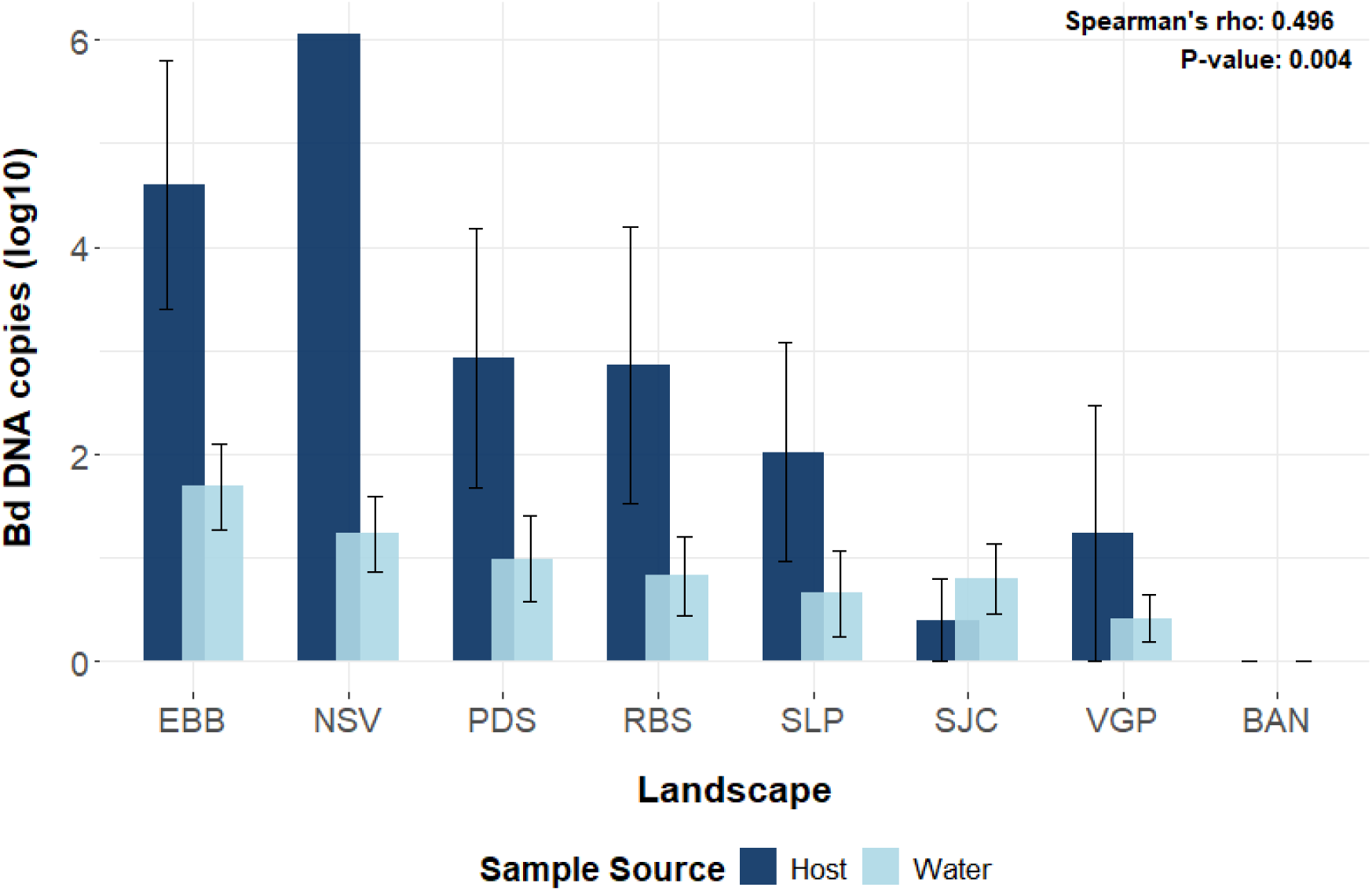
Log10-transformed *Bd* loads in amphibian hosts (total *Bd* DNA copies per frog, dark blue) and *Bd* concentration (*Bd* DNA copies/L, light blue) in aquatic samples collected from our eight focal study landscapes, all data were qPCR estimated. Landscapes are ordered from the most pristine (EBB) to the most impacted by habitat loss (BAN). Frog swabs from NSV were collected at a single location (mean = 6.06, se = 6.56). Landscape abbreviations: EBB, Estação Biológica de Boracéia; NSV, Núcleo de Santa Virgínia – Parque Estadual Serra do Mar; PDS, Pilar do Sul; RBS, Reserva Biológica Serra do Japi; SLP, São Luiz do Paraitinga; SJC, São José dos Campos; VGP, Vargem Grande Paulista; BAN, Bananal.

### 3.3. Landscape Integrity and *Bd* in the Environment

We found that the distribution of free-living *Bd* in water samples differs between streams in predominantly continuous and streams in heavily fragmented forests. Specifically, we found higher *Bd* DNA concentration (Wilcoxon rank-sum test: W = 323, *p* < 0.001) and occurrence (χ² = 10.025, d.f. = 1, *p* = 0.001) in stream water samples located in continuous forested landscapes compared to those from heavily fragmented forests (Figure 3).

**Figure 3.**
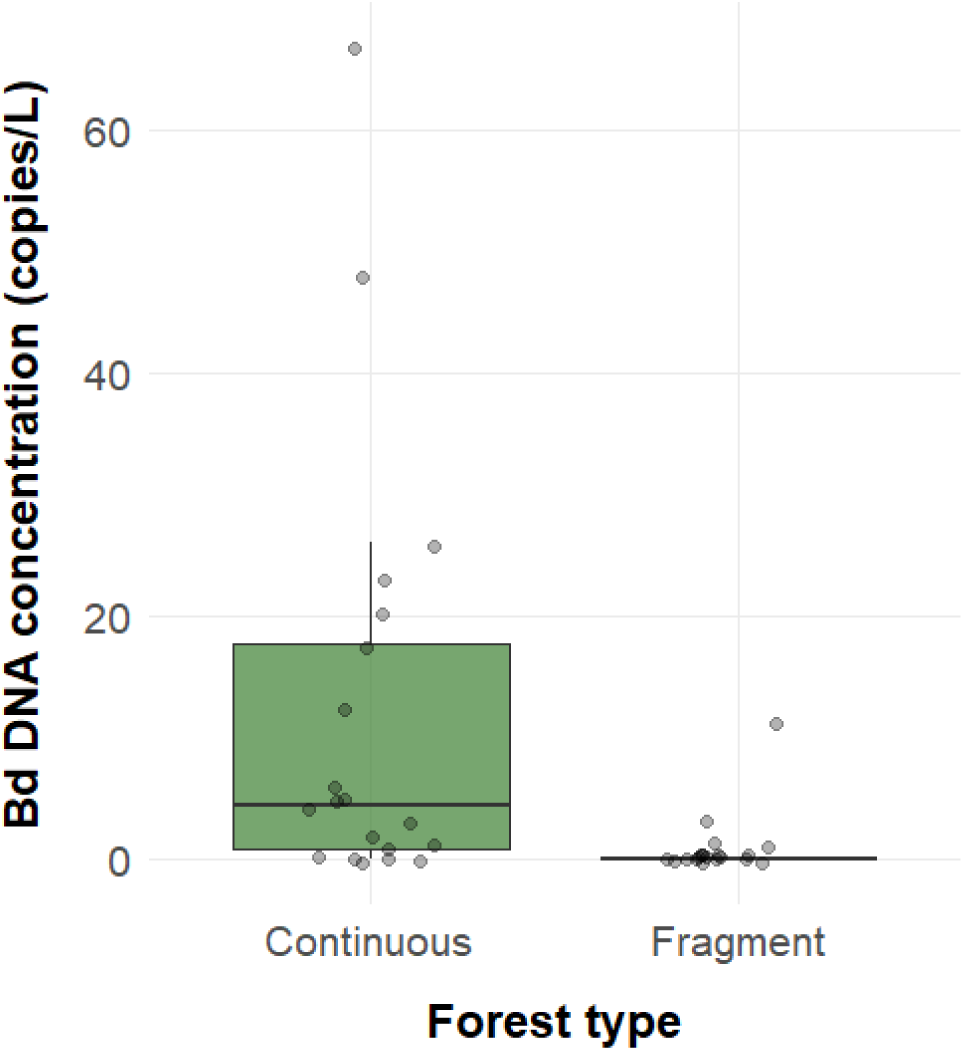
Distribution of free-living *Bd* in streams from predominantly continuous (Continuous) and heavily fragmented (Fragment) forests, as determined by dPCR (*Bd* DNA copies/L). Each data point represents an individual stream sample.

### 3.4. Landscape Integrity and Host-associated *Bd*

Our focal amphibian species exhibited distinct patterns of *Bd* infection load (Kruskal-Wallis = 82.057, d.f. = 3, p < 0.000). *Ischnocnema henselii* showed higher *Bd* load than the other species, while *Bd* was absent in *H. binotatus*, leading to its exclusion from downstream analyses (Figure SI 1). We observed species-specific effects of landscape integrity on *Bd* load within host populations. *Boana faber* exhibited higher *Bd* loads in continuous forests (Wilcoxon rank-sum test, W = 1249, p < 0.001). In contrast, neither *R. ornata* (Wilcoxon rank-sum test, W = 128, p = 0.668) nor *I. henselii* (Wilcoxon rank-sum test, W = 401, p = 0.094) exhibited a significant relationship with forest type in terms of *Bd* load.

The effect of landscape integrity on *Bd* occurrence also varied among species identity. *Ischnocnema henselii* showed a higher *Bd* prevalence (61%) in landscapes with high levels of habitat loss but showed similar patterns of *Bd* occurrence between continuous and fragmented forests (χ² < 0.0, d.f. = 1, p = 1). *Rhinella ornata* showed 20% prevalence in continuous forests, also with no significant difference in occurrence between forest types (χ² = 0.00, d.f. = 1, p = 1). However, *B. faber* exhibited significantly higher *Bd* occurrence in continuous forests (χ² = 14.1, d.f. = 1, p < 0.001; prevalence = 33%, Figure 4).

**Figure 4.**
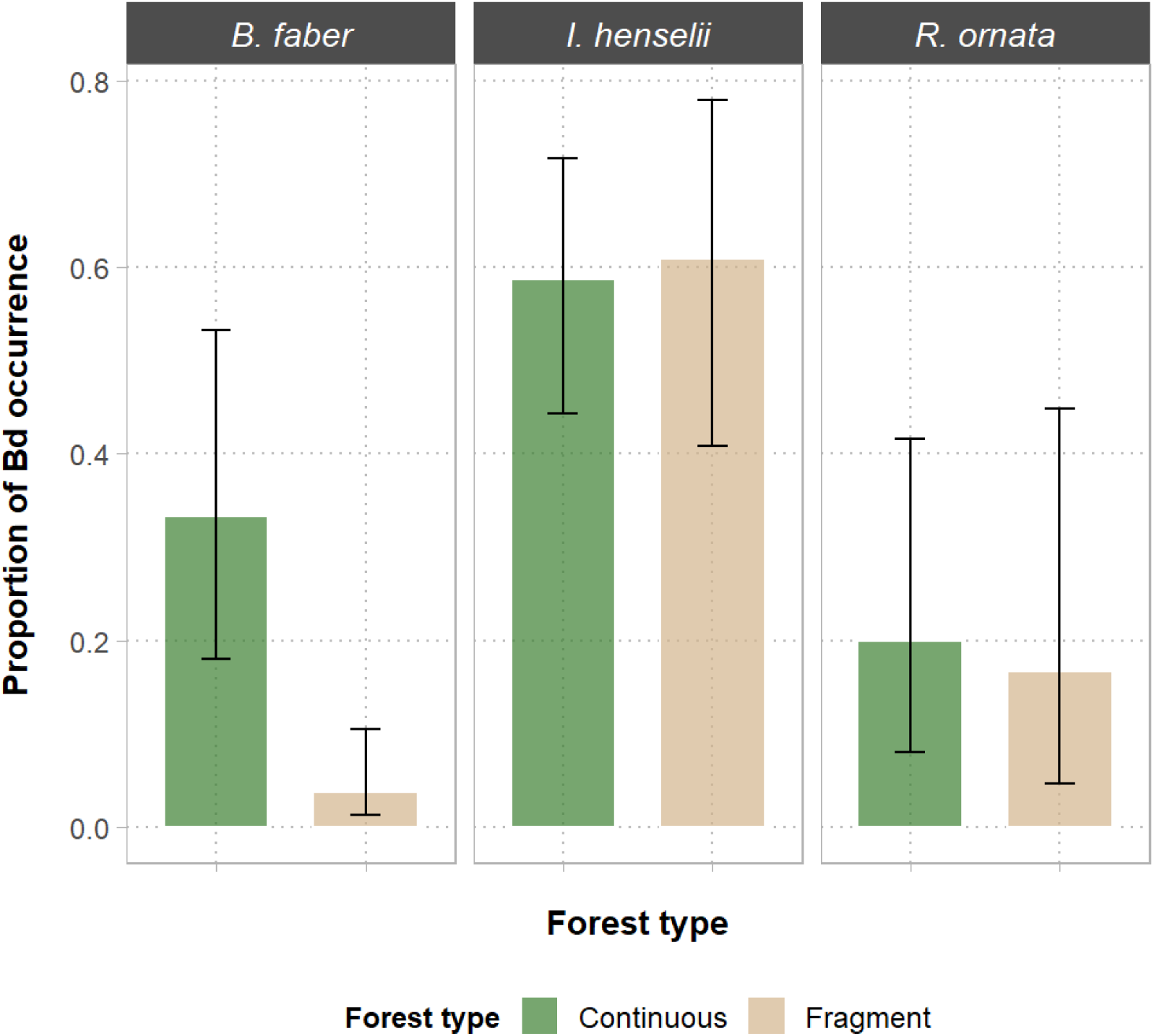
Bar chart illustrates *Bd* occurrence in three focal amphibian species: *Boana faber*, *Ischnocnema henselii* and *Rhinella ornata* in predominantly continuous (Continuous) and heavily fragmented (Fragment) forest landscapes. Error bars represent 95% binomial confidence intervals.

### 3.5. Influence of Habitat Structure – Water Samples

Our analysis identified key habitat structure metrics associated with variations in landscape integrity. Our results indicate that habitat structure markedly affects the distribution of *Bd* in water samples. At the landscape scale, forest cover emerged as the most substantial predictor of *Bd* concentration (Estimate = 1.728, *p =* 0.003) and probability of *Bd* occurrence (Estimate = 1.071, *p =* 0.006). Specifically, continuous and broader sections of natural forest cover were associated with elevated *Bd* concentrations (Figure 5a) and higher *Bd* occurrence within stream environments (Table 1). At the local scale, both *Bd* concentration (Estimate = −4.182, *p =* 0.004) and occurrence (Estimate = −3.466, *p =* 0.042) exhibited a strong negative relationship with habitat split (Table 1, Figure 5b), suggesting that increased habitat split is associated with a decrease in *Bd* distribution in streams.

**Figure 5.**
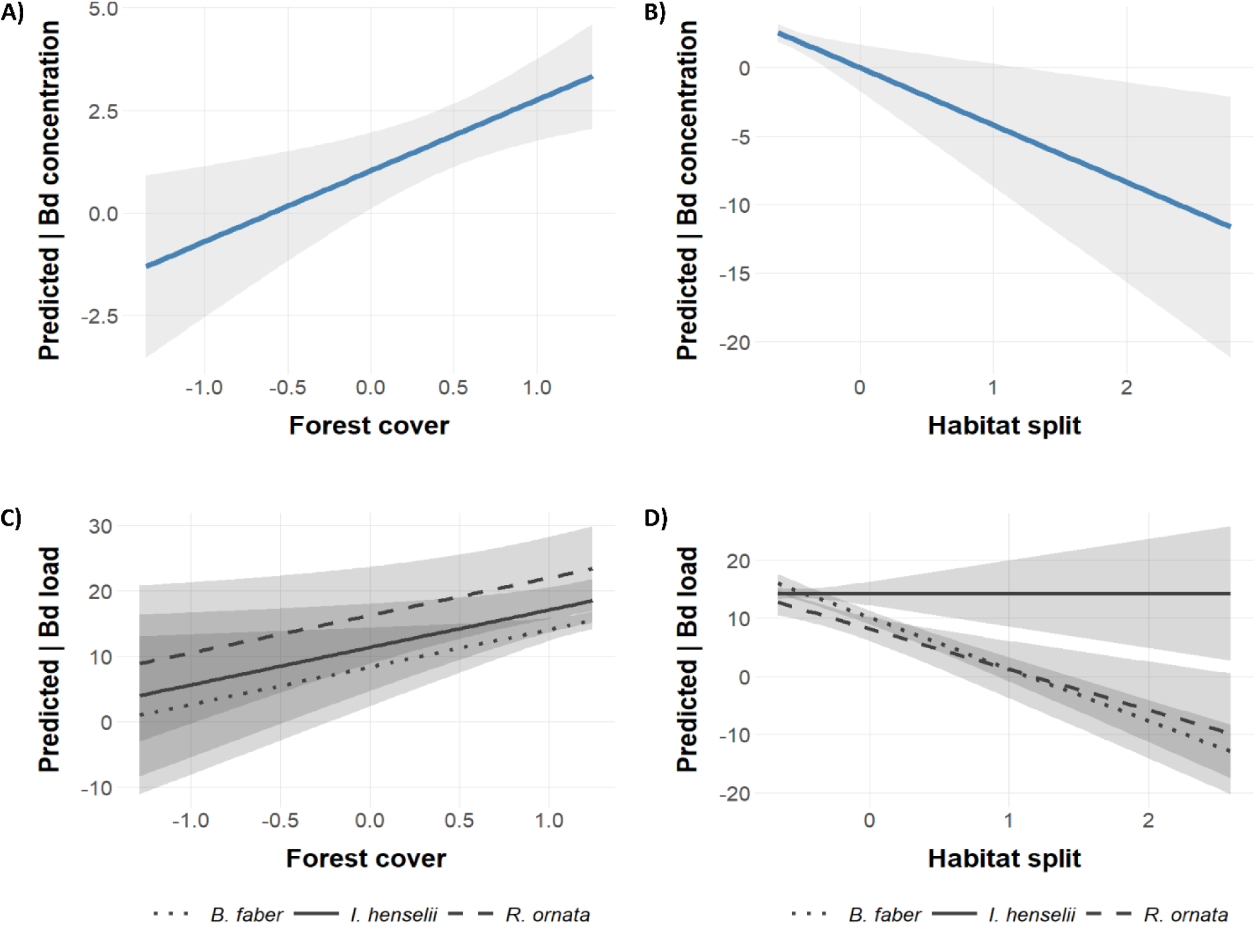
Predicted *Bd* DNA concentration (*Bd* DNA copies/L) in water bodies (A and B) and *Bd* infection load (*Bd* DNA copies per skin swab) in amphibian host species (C and D) across varying levels of forest cover and habitat fragmentation. All models were constructed using a negative binomial generalized linear mixed model (GLMM). Statistical analyses are detailed in Tables 1 and 2. The solid line represents the predicted relationship, while the shaded area indicates the 95% confidence interval around the predicted values.

**Table 1.** Most parsimonious Generalized Linear Mixed Models (GLMMs) for the distribution of *Bd* DNA across focal landscapes. *Bd* occurrence (presence/absence) was modeled using a binomial distribution, whereas *Bd* DNA concentration was modeled with a negative binomial distribution, both quantified via dPCR. The table presents model coefficients with ΔAIC ≤ 2 and their respective weights of evidence, indicating model support.

| **Landscape scale (10,000 m)** | Estimate | SD | z value | Pr ($>\vert z\vert $) |
| --- | --- | --- | --- | --- |
| *Bd* concentration; $\Delta AIC = 0.0$, $w_i = 0.762$ | | | | |
| Intercept | 1.094 | 0.422 | 2.593 | 0.010 |
| Forest cover | 1.728 | 0.589 | 2.935 | 0.003 |
| Connectivity | 0.218 | 0.541 | 0.402 | 0.687 |
| *Bd* occurrence; $\Delta AIC = 0.0$, $w_i = 0.619$ | | | | |
| Intercept | -0.136 | 0.359 | -0.379 | 0.705 |
| Forest cover | 1.071 | 0.387 | 2.765 | 0.006 |
| **Local scale (250 m)** | Estimate | SD | z value | Pr ($>\vert z\vert $) |
| *Bd* concentration; $\Delta AIC = 0.0$, $w_i = 0.731$ | | | | |
| Intercept | 0.114 | 0.896 | 0.127 | 0.899 |
| Percentage of local drainage | -0.641 | 0.387 | -1.658 | 0.097 |
| Habitat split | -4.182 | 1.456 | -2.873 | 0.004 |
| *Bd* occurrence; $\Delta AIC = 0.5$, $w_i = 0.344$ | | | | |
| Intercept | -1.243 | 0.932 | -1.333 | 0.182 |
| Habitat split | -3.466 | 1.703 | -2.036 | 0.042 |

**Table 2.** Most parsimonious Generalized Linear Mixed Models (GLMMs) for the distribution of *Bd* infecting amphibian skin across focal landscapes. *Bd* occurrence (presence and absence) was modeled using a binomial distribution, whereas the *Bd* infection load was modeled with a negative binomial distribution, both quantified through qPCR. The table presents the coefficients for models with ΔAIC ≤ 2 and their respective weights of evidence, indicating model support.

| **Landscape scale (10,000 m)** | Estimate | SD | z value | Pr ($>\vert z\vert $) |
| --- | --- | --- | --- | --- |
| *Bd* load; $\Delta AIC = 0.0$, $w_i = 0.612$ | | | | |
| Intercept | 6.656 | 2.542 | 2.618 | 0.008 |
| Forest cover | 5.753 | 2.438 | 2.36 | 0.018 |
| *Bd* occurrence; $\Delta AIC = 0.0$, $w_i = 0.908$ | | | | |
| Intercept | -2.113 | 0.572 | -3.692 | <0.001 |
| Drainage percentage | 1.594 | 0.607 | 2.627 | 0.009 |
| Drainage percentage* Frog species: *I. henselii* | -1.855 | 0.471 | -3.94 | <0.001 |
| Drainage percentage* Frog species: *R. ornata* | -1.391 | 0.554 | -2.51 | 0.012 |
| **Local scale (250 m)** | Estimate | SD | z value | Pr ($>\vert z\vert $) |
| *Bd* load; $\Delta AIC = 0.0$, $w_i = 0.364$ | | | | |
| Intercept | 10.168 | 0.602 | 16.883 | < 2e-16 |
| Habitat split | -8.908 | 0.883 | -10.091 | < 2e-16 |
| Habitat split * Frog species: *I. henselii* | 8.909 | 2.097 | 4.249 | <0.001 |
| Habitat split * Frog species: *R. ornata* | 1.934 | 2.013 | 0.961 | 0.337 |
| *Bd* occurrence; $\Delta AIC = 0.0$, $w_i = 1$ | | | | |
| Intercept | -2.42 | 0.447 | -5.412 | <0.001 |
| Forest cover | 1.492 | 0.393 | 3.794 | <0.001 |
| Forest cover * Frog species: *I. henselii* | -2.963 | 0.534 | -5.548 | <0.001 |
| Forest cover * Frog species: *R. ornata* | -1.482 | 0.584 | -2.535 | 0.011 |

### 3.6. Influence of Habitat Structure – Host Skin Samples

In agreement with the patterns observed for *Bd* distribution in aquatic environments, our models most strongly supported for explaining *Bd* distribution within host populations incorporated both forest cover and habitat split, in addition to the percentage of drainage in riparian protected areas (Table 2). At the landscape scale, *Bd* infection load was positively associated with forest cover (*Estimate* = 5.753, *p* = 0.018; Figure 5c), and the percentage of drainage emerged as the strongest predictor of *Bd* occurrence (*Estimate* = 1.594, *p* = 0.009). Interestingly, the positive relationship between the percentage of drainage and *Bd* occurrence appeared weaker or even reversed in *I. henselii* (*Estimate* = −1.855, p < 0.001) and *R. ornata* (*Estimate* = −1.391, *p* = 0.012, Figure SI 2).

At the local scale, increased habitat split was associated with reduced *Bd* infection loads in amphibian host species (*Estimate* = −8.908, *p* < 0.001). An interaction effect between habitat split and frog species was also identified (Table 2). The influence of habitat split differed for *I. henselii* compared to *B. faber,* which is the reference species in the models (*Estimate* = 8.909, *p* < 0.001), suggesting that the effect of reduced habitat split on *Bd* loads is reduced or reversed for *I. henselii* (Figure 5d). A model incorporating the interaction between forest cover (local scale) and amphibian species provided the best explanation for *Bd* occurrence (Table 2). This model indicated a positive association between forest cover and the likelihood of *Bd* occurrence (*Estimate* = 1.492, *p* < 0.001), suggesting enhanced detectability in areas with greater forest cover.

The combined effects revealed that the influence of forest cover on *Bd* occurrence varies by frog species. The negative interaction terms for *I. henselii* (*Estimate* = −2.963, *p* < 0.001) and *R. ornata* (*Estimate* = −1.482, *p* = 0.011) imply that the positive relationship between forest cover and *Bd* occurrence is reversed and weakened for these species, respectively (Figure SI 3).

### 3.7. *Bd* genotypes

Analysis of our focal species revealed two *Bd* lineages across the study region (Table SI 3). The globally distributed *Bd*-GPL lineage was dominant, while the enzootic *Bd*-Asia-2/Brazil lineage was detected only at two of our focal study landscapes: Estação Biológica de Boracéia and Pilar do Sul. Notably, co-infection with both lineages was confirmed for *B. faber* at Estação Biológica de Boracéia, although the presence of a hybrid lineage could not be confirmed or ruled out.

## 4. Discussion

Our study reveals a strong link between the distribution of *Bd* in aquatic environments and within amphibian host populations, a relationship that was strongly associated with habitat features at both the landscape and local scales. We found that the distribution patterns of *Bd* in the environment mirrored those found in host populations. More specifically, higher concentrations and occurrence of *Bd* in water bodies were also noted in more pristine habitats, as predicted by previous studies using only amphibian skin swabs (Becker et al., 2012; Becker and Zamudio, 2011). Our findings underscore the importance of incorporating environmental sampling into field surveys and *Bd* monitoring programs, as this approach provides valuable information on *Bd* persistence, dynamics, and potential transmission at the environmental level.

Our results from the comparison of forest categories were validated by findings from spatial modeling, in that higher forest cover and reduced habitat split, found in pristine continuous forests, were associated with increased *Bd* occurrence and concentration in water bodies. The structural complexity of well-preserved forests buffers against temperature increases and promotes humidity retention (De Frenne et al., 2019). Therefore, lower temperatures of these habitats may be linked to a higher prevalence and infection intensity in amphibians (Gass et al., 2024; Herczeg et al., 2023; Piotrowski et al., 2004); while higher humidity levels may promote a decreased *Bd* infection loads and improved amphibian survival (Gass et al., 2024). Additionally, habitat connectivity and water availability are likely to create more suitable conditions for *Bd* persistence and transmission through environmental sources, particularly given its aquatic life stage. Other environmental factors associated with aquatic ecosystems, such as temperature and pH (Piotrowski et al., 2004), salinity (Clulow et al., 2018; Stockwell et al., 2015), and exposure to contaminants (Rohr et al., 2017), also have been shown to influence the survival and infectivity of *Bd* zoospores. While most *Bd* research focuses on host-based approaches (Mosher et al., 2018), our study demonstrates the effect of the habitat preservation on the occurrence and concentration of free-living *Bd*, at the environment level, advancing our understanding of its spatial distribution outside amphibian hosts.

In amphibian hosts, we found that higher natural vegetation cover, drainage percentage, and habitat connectivity favor higher *Bd* occurrence and infection load, mirroring the pattern observed in aquatic environments. These findings corroborate previous research that found higher *Bd* loads and prevalence in amphibians from pristine, continuous forest environments (Becker et al., 2012; Becker and Zamudio, 2011). A possible complementary explanation for the higher occurrence and concentration of environmental *Bd* in less-disturbed habitat is that these forests also have a greater number of hosts, due to the more favorable conditions for their survival. It is important to note that, based on this study, we cannot determine whether the low environmental concentration of *Bd* in more opened canopy areas results from a reduced prevalence and load in amphibians; or if the lower loads in amphibians are a consequence of the low environmental concentrations and occurrence in the more deforested habitats. Further, we cannot conclude that a decreased concentration of *Bd* in aquatic environments, potentially influenced by environmental factors such as temperature, pH, salinity, and contaminants (Clulow et al., 2018; Piotrowski et al., 2004; Rohr et al., 2017; Stockwell et al., 2015), contribute to a decrease in *Bd* prevalence and load within amphibian populations.

While well-preserved forests have higher *Bd* occurrences, the interaction with amphibian species highlights the complex interplay of environmental and host factors in driving *Bd* distribution. This pattern is in agreement with previous studies that observed differences in *Bd* infection patterns among amphibian species, indicating that host physiological, ecological, evolutionary history, and even microbiome profiles are involved in variations of host susceptibility (Bates et al., 2022; Longo et al., 2023; Voyles et al., 2018). Species-specific traits can affect *Bd* exposure and infection rates, with some Bd-tolerant host species acting as asymptomatic carriers that amplify disease transmission, while other species being highly susceptible (Longo et al., 2023). Given that intrinsic physiological processes can themselves be modulated by environmental factors, variations in host-*Bd* susceptibility may reflect a dynamic interplay between endogenous capacities and environmental modulation, rather than being solely the product of fixed, environment-independent traits.

*Ischnocnema henselii*, in particular, displays weaker interaction with habitat split locally and drainage percentage regionally, potentially indicating limited reliance of this species on aquatic environments. As a species with direct terrestrial development, *I. henselii* may be more strongly influenced by factors related to the terrestrial environment, such as the depth of the leaf litter layer, soil type, and clay content (Menin et al., 2007; Oliveira et al., 2013). In contrast, amphibian species with aquatic larval development, such as *B. faber* and *R. ornata*, could be more susceptible to factors related to their aquatic reproductive sites (Menin et al., 2011; Parris and McCarthy, 1999). Despite sharing direct terrestrial development and ecological similarities with *I. henselii* (Haddad et al., 2013), the lack of infection observed in *H. binotatus* suggests that endogenous host factors, along with environmental modulation, play a key role in its response to infection. Our finding corroborates previous research reporting similar patterns in *H. binotatus* (Martins et al., 2022), suggesting that differences in bacteriome structure may contribute to varying susceptibility/resistance to chytridiomycosis (Bates et al., 2022). When examining the effects of habitat split on host *Bd* infection, it is crucial to consider host movement/migration patterns during the breeding season, between upland forest fragments and lowland streams that are often located in open and disturbed environments (Becker et al., 2010). Because breeding migrations shift direction towards the end of the rainy season (most frog species move back to upland forests), exposure to *Bd* and the development of acquired resistance to waterborne pathogens is also expected to shift temporally (Becker et al., 2023). Therefore, individual frogs sampled later in the breeding season (January to March), as in our study, may show different *Bd*-environment associations compared to those that might have been observed if sampling had occurred at the onset of the breeding migration. Patterns of environmental *Bd* distribution might also not align with *Bd* infection patterns in host populations throughout the year. Therefore, future studies should consider environmental, seasonal, and behavioral and physiological aspects of the studied amphibian species.

While *Bd* monitoring depends on the techniques employed, we have demonstrated an equivalence between digital PCR (dPCR) and quantitative PCR (qPCR) in detecting and quantifying habitat-level *Bd* when using our high-capacity filtration method. The challenges in detecting environmental *Bd* are likely driven by the low volume of water filtration. Even without a concrete link between water volume and *Bd* quantification, our successful environmental *Bd* detection is attributed to the large filtration volumes (10 - 255 L), and these results are in accordance with proposed methods by Brannelly et al. (2020). Method selection should also consider factors beyond sample volume, including the strengths and weaknesses of each PCR-based technique, DNA extraction methods, sample type, study aims, detection time, and cost (Brannelly et al., 2020; Zhang et al., 2024). For example, dPCR is more sensitive than qPCR, has increased precision, and allows for the absolute quantification of rare targets without the need for a standard as per the manufacturer (ThermoFisher Scientific). Effective monitoring levering newer molecular diagnostic technology is essential for predicting and mitigating local and regional epizootics, as well as for better estimating host-pathogen dynamics.

Our study demonstrated an association between environmental and host-associated *Bd*, which remained consistent across a range of habitat quality. Our findings highlight that environmental degradation is associated with a decrease in environmental *Bd* concentration (or host infection loads) and *Bd* occurrence in aquatic environments and host populations. Predicting wildlife disease outbreaks presents a formidable challenge due to their dependence on fluctuations in host, pathogen, and environmental factors, along with the complex interactions among these elements. Moreover, the efficacy of disease management strategies is contingent upon the availability of relevant data. Here we propose a novel method with large-volume filtration and dual dPCR/qPCR validation for detecting and quantifying the primary pathogen currently affecting amphibians, which can inform *Bd* modeling in amphibian populations but without the need for direct sampling of amphibian hosts.

## CRediT authorship contribution statement

A. B. Assis: Writing – Conceptualization, Data curation, Formal analysis, Funding acquisition, Investigation, Methodology, Project administration, Resources, Validation, Visualization, Writing – original draft, Writing – review & editing. D. Rodriguez: Conceptualization, Methodology, Resources, Validation, Writing – review & editing. W. J. Neely: Formal analysis, Investigation, Validation, Writing – review & editing. C. M. Lopes: Investigation, Methodology, Writing – review & editing. P. Prist: Investigation, Validation, Writing – review & editing. C. A. Navas: Resources, Writing – review & editing. R. A. Martins: Investigation. C. F. B. Haddad: Supervision, Conceptualization, Funding acquisition, Investigation, Resources, Validation, Visualization, Writing – review & editing. C. G. Becker: Supervision, Conceptualization, Formal analysis, Funding acquisition, Investigation, Methodology, Resources, Validation, Visualization, Writing – review & editing.

## Acknowledgment

The authors express their sincere gratitude for all the financial support that made this project possible. We also deeply appreciate the cooperation of the landowners who generously granted access to their properties for sample collection. Special thanks to Vagner Alberto for his laboratory assistance, Lis Batista Primo for her contributions to DNA extractions, and Luis F. Montes for his support in fieldwork.

## Compliance with ethical standards

### Ethical approval

All work was conducted under appropriate permits. Instituto Chico Mendes – SISBIO #74576-5; SISGEN #AC2F7B2/ #R3B282B; Governo do Estado de São Paulo / Secretaria de Infraestrutura e Meio Ambiente / Instituto de Pesquisas Ambientais – IPA #IF.003838/2020-26.

