## Supplementary Information (SI) for "Environmental Detection of the Amphibian Chytrid Fungus in Water Bodies Predicts Host Infection Along a Deforestation Gradient"

**Table SI 1. Summary of the total samples collected across our focal landscapes.** Each row represents a specific landscape, detailing the number of water samples and host samples collected.

| Landscape | Landscape type | Sites | Water | Host |
|---|---|---|---|---|
| Estação Biológica de Boracéia | Continuous | 5 | 5 | 61 |
| Núcleo de Santa Virgínia – PESM | Continuous | 5 | 5 | 10 |
| Reserva Biológica Serra do Japi | Continuous | 5 | 5 | 15 |
| Pilar do Sul | Continuous | 5 | 5 | 22 |
| Vargem Grande Paulista | Fragmented | 5 | 5 | 50 |
| São Luiz do Paraitinga | Fragmented | 5 | 5 | 38 |
| Bananal | Fragmented | 5 | 5 | 48 |
| São José dos Campos | Fragmented | 5 | 5 | 59 |

### LANDCOVER METRICS

Spatial data at the landscape and sampling site levels, including hydrological features were obtained from the land use land cover mapping of the Brazilian Foundation for Sustainable Development (Fundação Brasileira Para o Desenvolvimento Sustentável). This mapping was generated using RapidEye satellite imagery at a 1:10,000 scale, having a spatial resolution of 5 x 5 meters and corresponded to the year 2017 (Fundação Brasileira Para o Desenvolvimento Sustentável, 2020).

The land use / land cover mapping encompasses six landscape covers including both natural habitats as well as anthropogenic disturbances such as agriculture, silviculture, and urbanization. Riparian protected areas are also mapped through the Brazilian Foundation for Sustainable Development and were integrated into the land use land cover mapping. Hydrological features included drainage networks comprising streams and ponds, and to ensure accuracy, small permanent water bodies not captured by the Brazilian Foundation for Sustainable Development mapping were mapped by visual classification in Google Earth and in ArcGis and integrated into this dataset. The final land use land cover mapping used combined the land use cover, hydrological features and the riparian protected area maps.

From this dataset, we derived eighteen landscape metrics measured at the landscape (10 km radius) and local scales (250 m radius) and considering both landscape cover and configuration: percentage of forest cover, percentage of drainage, edge density, habitat split (*i.e.*, spatial discontinuity between natural forest vegetation and aquatic breeding sites such as streams and ponds) and connectivity, see Table SI 2.

Percentage of forest cover, drainage, water mass and riparian protected areas correspond to the percentage of the landscape comprised of this land use type. It was calculated considering the sum of the area of all fragments corresponding to this land use type, divided by the total landscape area and multiplies by 100 to convert to a percentage. Number of forest fragments corresponds to the number of patches of this land use type. Drainage density and forest edge density correspond to the sum of the lengths of all edge segments in the landscape (corresponding to this land use type), divided by the total landscape area, multiplied by 10,000, to convert to hectares. Finally, euclidean nearest neighbor distance corresponds to the distance to the nearest neighbouring patch of the same type, based on shortest edge-to-edge distance. By examining variation in habitat split among forest fragments with comparable sizes and edge densities, we isolated the effects of forest fragmentation from those of habitat split.

**Table SI 2**. Landscape variables evaluated.

| **Variable** | **Spatial scale (radius)** |
|---|---|
| **Forest** | |
| Percentage of forest cover | 250 m / 10,000 m |
| Percentage of riparian protected area | 10,000 m |
| Percentage of forest fragmente | 10,000 m |
| Number of forest fragments inside 10 km buffer | 10,000 m |
| Fragment size (area in m2) | 250 m |
| **Drainage** | |
| Drainage density (meters/hectares) | 250 m / 10,000 m |
| Drainage percentagem | 250 m / 10,000 m |
| Number of drainage | 250 m |
| Water mass percentagem | 10,000 m |
| Drainage percentage in riparian protected area | 10,000 m |
| Drainage density in riparian protected areas | 10,000 m |
| **Edge** | |
| Forest edge density (forest fragments and riparian protected areas)* | 250 m / 10,000 m |
| Edge density of forest fragments * | 250 m / 10,000 m |
| Edge density of riparian protected areas* | 250 m / 10,000 m |
| **Habitat split** | |
| Habitat split - ponds** | 250 m |
| Habitat split - streams** | 250 m |
| **Connectivity** | |
| Euclidean nearest neighbor distance - fragments *** | 10,000 m |
| Euclidean nearest neighbor distance - riparian protected areas *** | 10,000 m |

* Determined by summing the lengths (in meters) of all edge segments within a site, dividing by the total landscape area (in $m^2$), and multiplying by 10,000 to convert the ratio to meters per hectare

** The average minimum distance (north, south, east, and west) between the edge of natural habitat and the nearest pond or stream.

*** The distance from a patch to a neighboring patch, based on the nearest cell center-to-cell center. Here we are using the mean distance of all patches inside the buffer; Distances close to zero = more aggregated; Unit (meters)

**Table SI 3.** Distribution of *Batrachochytrium dendrobatidis* lineages across Brazilian landscapes in amphibian host species**.**

| **Landscape** | **Genotype** | **Host** |
|---|---|---|
| Bananal | *Bd*-absent | - |
| São Luiz do Paraitinga | *Bd*-GPL | *I. henselii* |
| Núcleo de Santa Virgínia | *Bd*-GPL | *I. henselii* |
| Estação Biológica de Boracéia | *Bd*-GPL / Bd-Asia-2/Brazil | *B. faber* / *R. ornata* / *I. henselii* |
| São José dos Campos | Not applicable | - |
| Reserva Biológica Serra do Japi | *Bd*-GPL | *R. ornata* / *I. henselii* |
| Vargem Grande Paulista | *Bd*-GPL | *I. henselii* |
| Pilar do Sul | *Bd*-GPL / Bd-Asia-2/Brazil | *B. faber* / *I. henselii* |

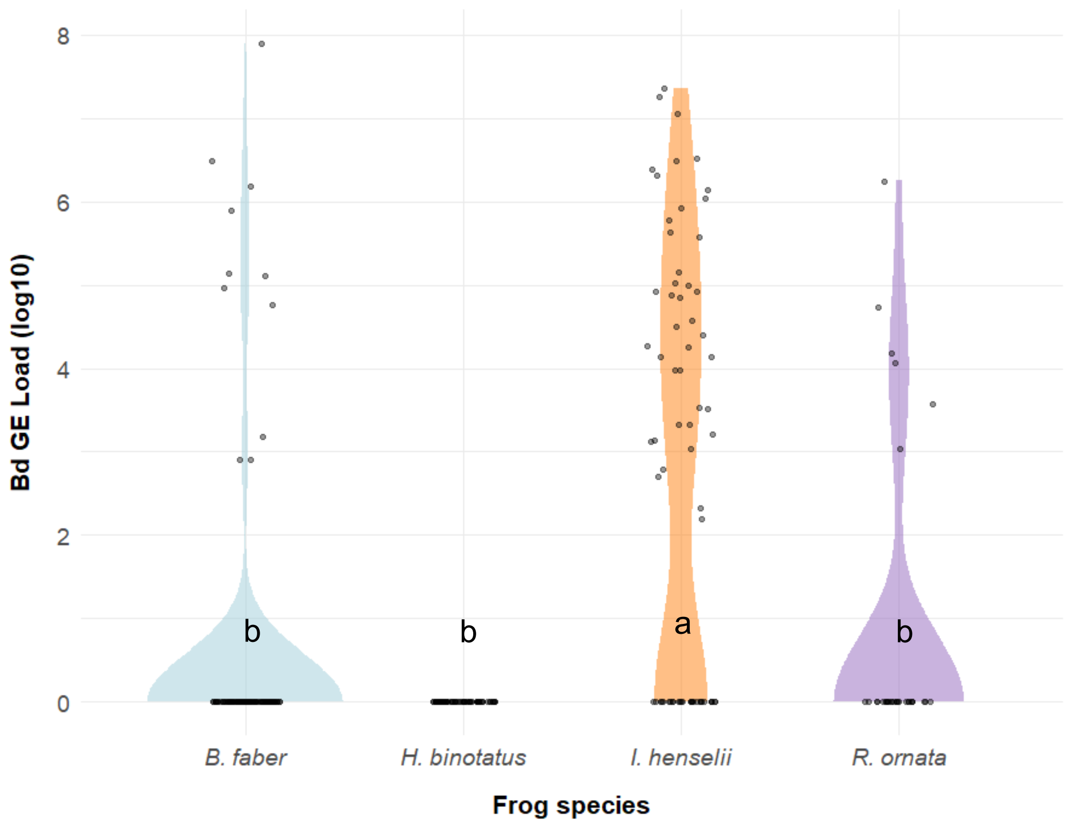


**Figure SI 1**. Violin plot of *Bd* load across the focal amphibian species. A Kruskal-Wallis's test indicates species' differences in *Bd* GE loads ($\chi^2 = 82.06$, df = 3, $P < 0.001$). Pairwise comparisons were performed using Dunn's test with Bonferroni correction. Different lowercase letters indicate statistically significant differences in *Bd* GE load among species at $\alpha = 0.05$, as determined by post hoc.

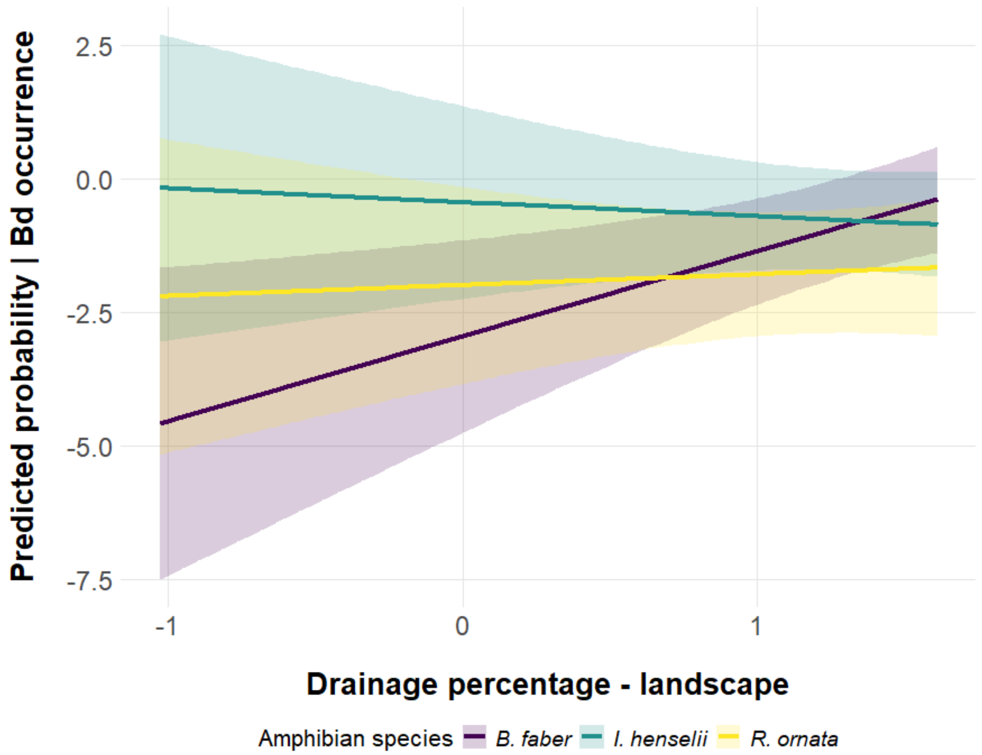


**Figure SI 2**. Predicted probability of *Bd* detection across varying drainage percentage – at the landscape scale – for three amphibian species. A binomial generalized linear mixed model (GLMM) was employed, including a random intercept for landscape to account for spatial autocorrelation and interaction terms to assess species-specific responses to drainage. Landscape-level drainage percentage had a positive effect on the probability of *Bd* detection ($p = 0.009$). Significant interaction ($p < 0.0001$ for *B. faber* and *I. henselii*, and $p = 0.012$ for *R. ornata*) were observed. Lines represent model predictions with 95% confidence intervals (shaded areas).

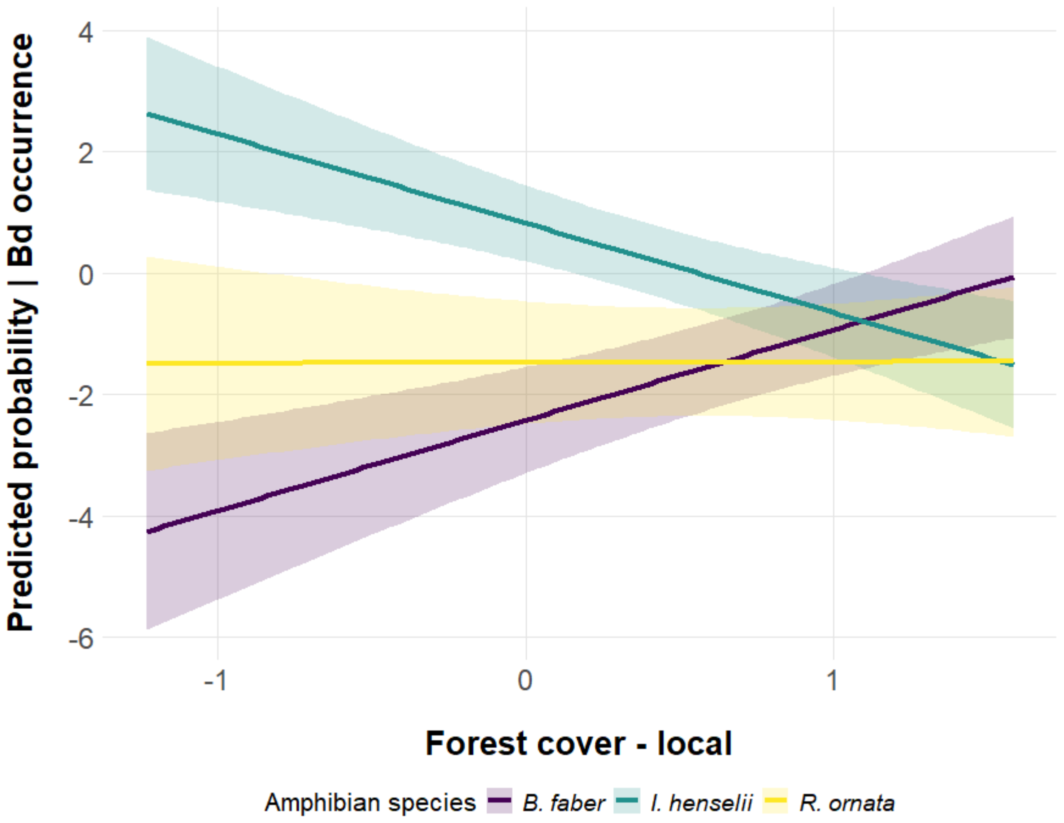


**Figure SI 3**. Predicted probability of Bd detection across varying levels of forest cover - at the local scale - for three amphibian species. A binomial GLMM with a random effect for landscape and an interaction term between forest cover and species was used. This model revealed a positive association between forest cover and Bd detection probability (Estimate = 1.492; $p < 0.001$). The model also indicates significant interactions between forest cover and species ($p < 0.001$ for *B. faber* and *I. henselii*; and $p = 0.011$ for *R. ornata*). Model predictions with 95% confidence intervals are presented (lines and shaded areas, respectively).
